# CodAn: predictive models for the characterization of mRNA transcripts in Eukaryotes

**DOI:** 10.1101/794107

**Authors:** Pedro G Nachtigall, Andre Y Kashiwabara, Alan M Durham

## Abstract

Characterization of the coding sequences (CDSs) is an essential step on transcriptome annotation. Incorrect characterization of CDSs can lead to the prediction of non-existent proteins that can eventually compromise knowledge if databases are populated with similar incorrect predictions made in different genomes. Even though some recent methods have succeeded in correctly prediction of the stop codon position in strand-specific sequences, prediction of the complete CDS is still far from a gold standard. More importantly, prediction in strand-blind sequences and in partial sequences is deficient, presenting very low accuracy. Here, we present CodAn, a new computational approach to predict CDS and UTR, that significantly pushes the boundaries of CDS prediction in strand-blind and in partial sequences, increases strand-specific full-CDS predictions and matches or surpasses gold-standard results in strand-specific stop codon predictions. CodAn is freely available for download at https://github.com/pedronachtigall/CodAn.

## Introduction

The complete characterization of sequences resulting from a transcriptome assembly is an important step to understand the profile of genes expressed in the sample [3]. The coding regions (CDS) of the transcripts represent the functional part of the protein-coding genes, which correlates to the biological function of that gene [16]. Also, the untranslated regions (UTRs) are considered crucial to understanding the genetic regulatory networks involved in specific biological pathways [20, 7, 21]. UTRs have been shown to be major components on post-transcriptional regulation of gene expression (reviewed by [1]). UTRs are responsible to regulate mRNA stability, export, cellular localization, and translation efficiency, which influence directly the final amount of protein (reviewed by [23]). Moreover, the complex pattern of UTR regulation is strongly associated with embryogenesis, cellular diversity and diseases [27, 7, 33]. The correct characterization of the UTR and CDS landscape is, therefore, an essential initial step in correctly identifying regulatory elements that can determine the final protein output.

Currently, there are several computational tools to detect the CDSs and UTRs of transcripts. Some of these tools focus on characterizing the CDS [19, 6, 29] and others in characterizing the UTR regions [5, 13, 14, 9, 32]. Additionally, some widely used machine learning approaches were developed to classify transcripts as protein-coding genes or non-coding genes [18, 34, 26, 11], but these methods are only classifiers and do not perform annotation of the coding sequences.

There are basically two strategies for the implementation of these predictors: similarity search, and *ab initio* predictors. Similarity-based methods [9, 31, 22] rely on the existence of curated proteins and are useful for genes that code for closely related curated proteins, but fail to characterize CDS for novel proteins. We can separate *ab initio* prediction methods in two categories: (i) pretrained methods [19, 25], that generally require curated sequences to estimate specific parameters or the use of the pre-computed parameters of the closely-related species available; and (ii) self-training methods, which detect putative long ORFs in the transcripts to train a prediction model specific to that set of sequences [6, 29, 2, 4, 10, 30].

The design of a computational tool that can be easily and automatically applicable to any species and to strand-specific, strand-blind or partial sequences, is necessary for a wide and confident characterization of CDS and UTR landscape in all novel transcriptome projects. Three previous approaches circumvent this problem with a self-training approach, where predictors first perform an expectation maximization (EM) interactive procedure to train the prediction model using the target data and can be appied to any organism. Of these, GeneMarkS-T [9] presents a performance closer to a gold standard in stop-codon prediction, with an average of more than 90% of correct predictions of stop codons [29]. However, as we will show below, performance decreases when considering full CDS prediction (i.e., correct start and stop codon identification), strand-blind prediction (where the orientation of the transcript is unknown) or partial sequence prediction, indicating the need for new approaches that can reliably characterize CDS in all sequencing scenarios.

Here we present CodAn, a new transcript characterization software that can be applied to any eukaryotic organism and that dramatically increases the current accuracy boundaries in partial and in strand-blind sequences, increases the accuracy in start codon predicion, and matches or surpassing gold-standard accuracy for stop codon predicion in strand-specific sequences. CodAn has four different probabilistic models for four groups of Eukaryotes: vertebrates, invertebrates, plants, and fungi. We show that with these pre-designed models, CodAn can perform highly confident predictions of the full CDS and UTR regions not only in strand-specific full transcript sequences but also in strand-blind and partial sequences in a rate far higher than other available software.

## Results and discussion

CodAn is a stand alone software that can be used to reliably predict the location of UTR and CDS regions in full or partial transcripts. CodAn uses two Generalized Hidden Markov Models, one for a full CDS and another for partial transcripts.

We compared CodAn’s performance against that of ESTScan [19], TransDecoder [6], Prodigal [10] and GeneMarkS-T [29] in 34 different organisms of four groups: vertebrates, invertebrates, plants, and fungi (Table 1). For each organism, the performance was measured in transcripts of eight different test sets: two sets of strand-specific full transcripts, one set of strand-blind full transcripts, three sets of partial transcripts (“No Start”, “No Stop”, “No Start & No Stop”, and two different negative sets (3’UTR partial transcripts and ncRNA transcripts).

**Table 1:**
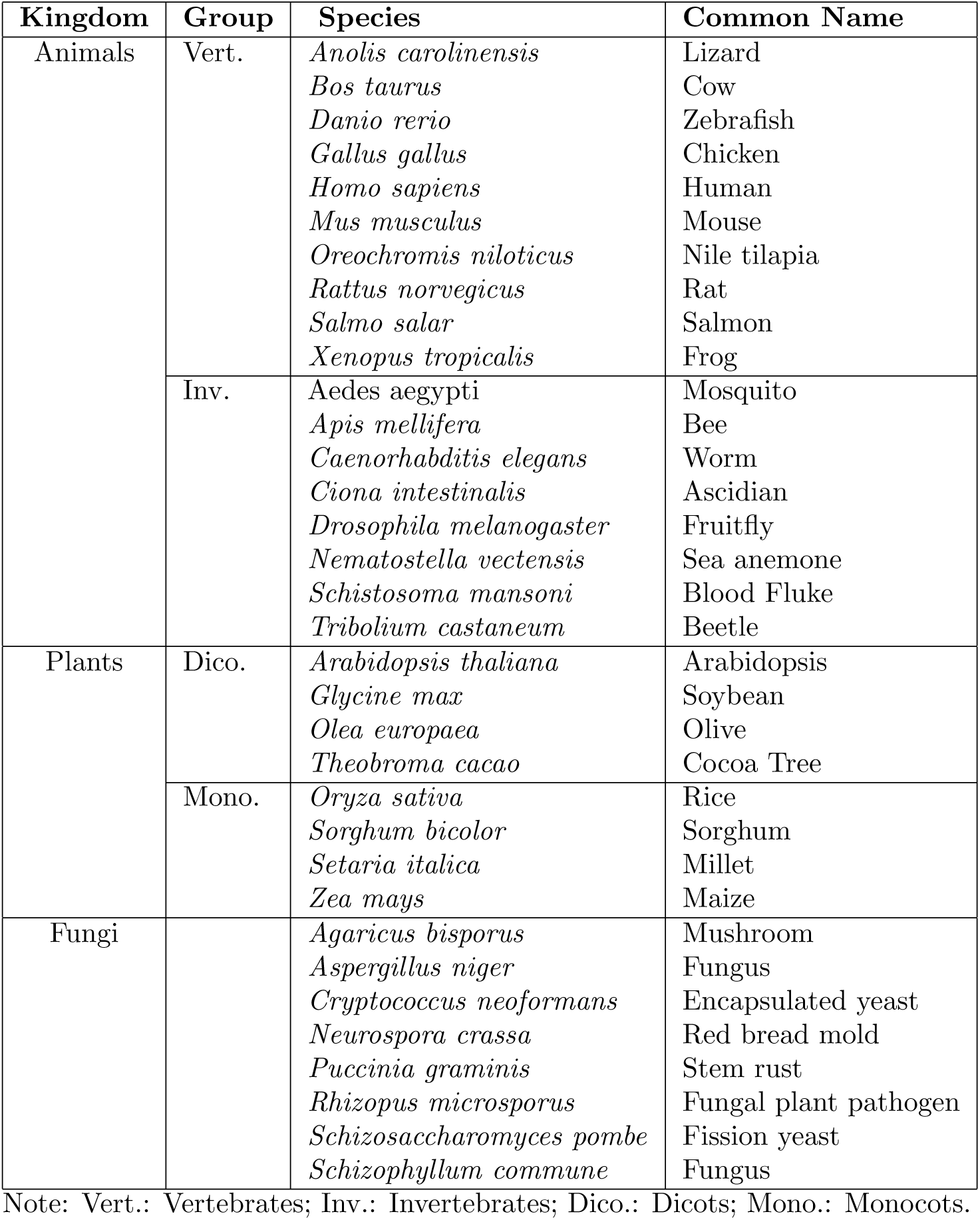
Species with validated annotations at RefSeq and used in the present study.

### Prediction accuracy assessment: full transcripts

For strand-specific stope codon prediction, CodAn presents a higher performance in all 4 groups as we can see in Table 2. Average F1-scores for each category were all above 97%, constantly higher than other approaches. Low standard deviation values in all four organism groups (equal or lower than 0.01) indicate the robustness of the method. This performance is confirmed when examining a summary of the results for each species, as depicted in Figure 1A, which shows the values obtained for Precision, Sensitivity, and F1-score. For complete strand-specific sets, CodAn presented a higher performance for the majority of the organisms in all 4 categories (Tables S2 A and S2 B at Supplemental Table S2).

**Table 2:**
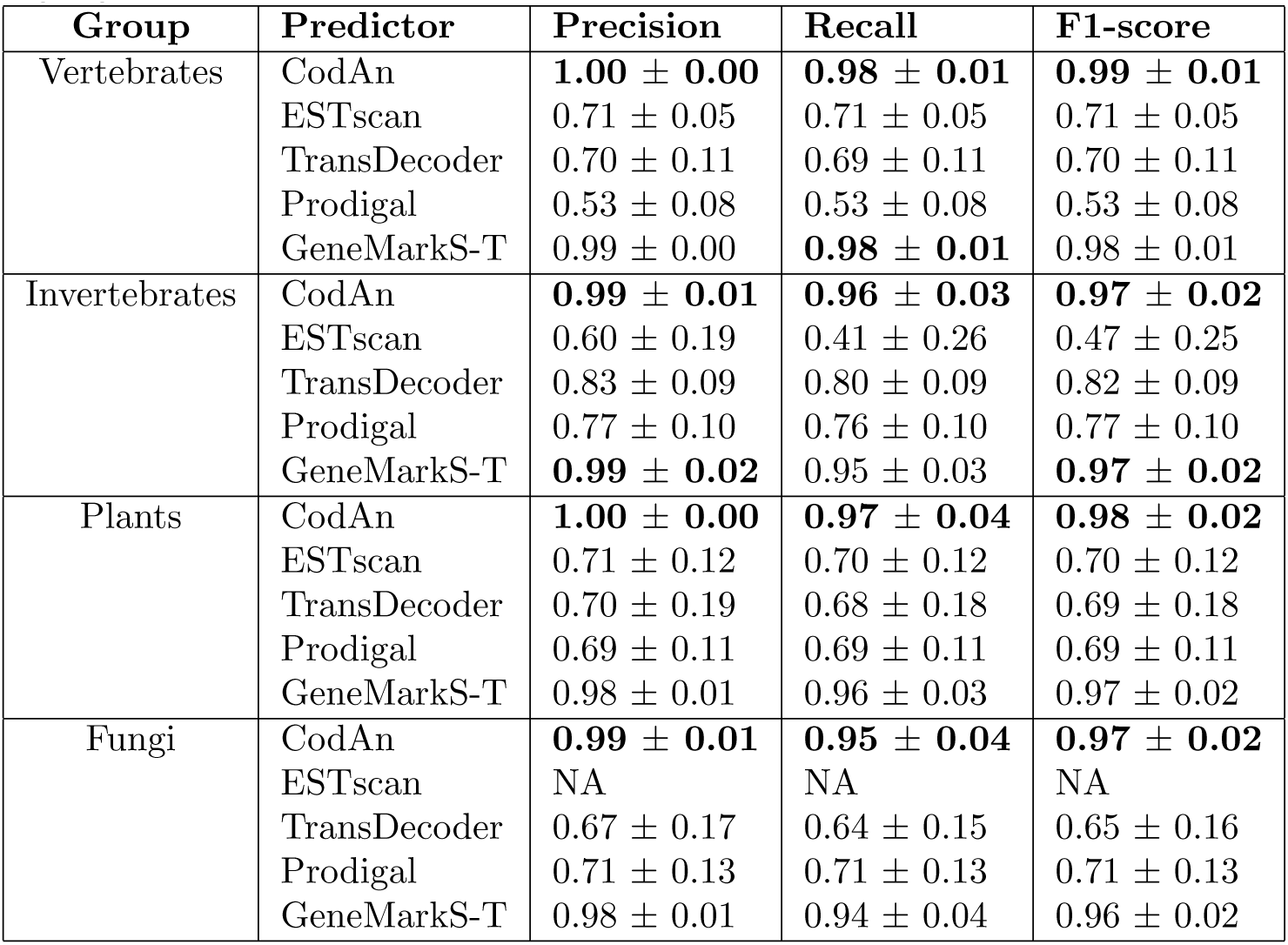
Average and standard deviation of precision, Recall and F1-score obtained by each tool in the strand-specific full transcript sets analyzed. Bold font highlight the higher value for each group.

| Group | Predictor | Precision | Recall | F1-score |
| --- | --- | --- | --- | --- |
| Vertebrates | CodAn | **1.00 ± 0.00** | **0.98 ± 0.01** | **0.99 ± 0.01** |
| | ESTscan | 0.71 ± 0.05 | 0.71 ± 0.05 | 0.71 ± 0.05 |
| | TransDecoder | 0.70 ± 0.11 | 0.69 ± 0.11 | 0.70 ± 0.11 |
| | Prodigal | 0.53 ± 0.08 | 0.53 ± 0.08 | 0.53 ± 0.08 |
| | GeneMarkS-T | 0.99 ± 0.00 | **0.98 ± 0.01** | 0.98 ± 0.01 |
| Invertebrates | CodAn | **0.99 ± 0.01** | **0.96 ± 0.03** | **0.97 ± 0.02** |
| | ESTscan | 0.60 ± 0.19 | 0.41 ± 0.26 | 0.47 ± 0.25 |
| | TransDecoder | 0.83 ± 0.09 | 0.80 ± 0.09 | 0.82 ± 0.09 |
| | Prodigal | 0.77 ± 0.10 | 0.76 ± 0.10 | 0.77 ± 0.10 |
| | GeneMarkS-T | **0.99 ± 0.02** | 0.95 ± 0.03 | **0.97 ± 0.02** |
| Plants | CodAn | **1.00 ± 0.00** | **0.97 ± 0.04** | **0.98 ± 0.02** |
| | ESTscan | 0.71 ± 0.12 | 0.70 ± 0.12 | 0.70 ± 0.12 |
| | TransDecoder | 0.70 ± 0.19 | 0.68 ± 0.18 | 0.69 ± 0.18 |
| | Prodigal | 0.69 ± 0.11 | 0.69 ± 0.11 | 0.69 ± 0.11 |
| | GeneMarkS-T | 0.98 ± 0.01 | 0.96 ± 0.03 | 0.97 ± 0.02 |
| Fungi | CodAn | **0.99 ± 0.01** | **0.95 ± 0.04** | **0.97 ± 0.02** |
|  | ESTscan | NA | NA | NA |
| | TransDecoder | 0.67 ± 0.17 | 0.64 ± 0.15 | 0.65 ± 0.16 |
| | Prodigal | 0.71 ± 0.13 | 0.71 ± 0.13 | 0.71 ± 0.13 |
| | GeneMarkS-T | 0.98 ± 0.01 | 0.94 ± 0.04 | 0.96 ± 0.02 |

**Figure 1:**
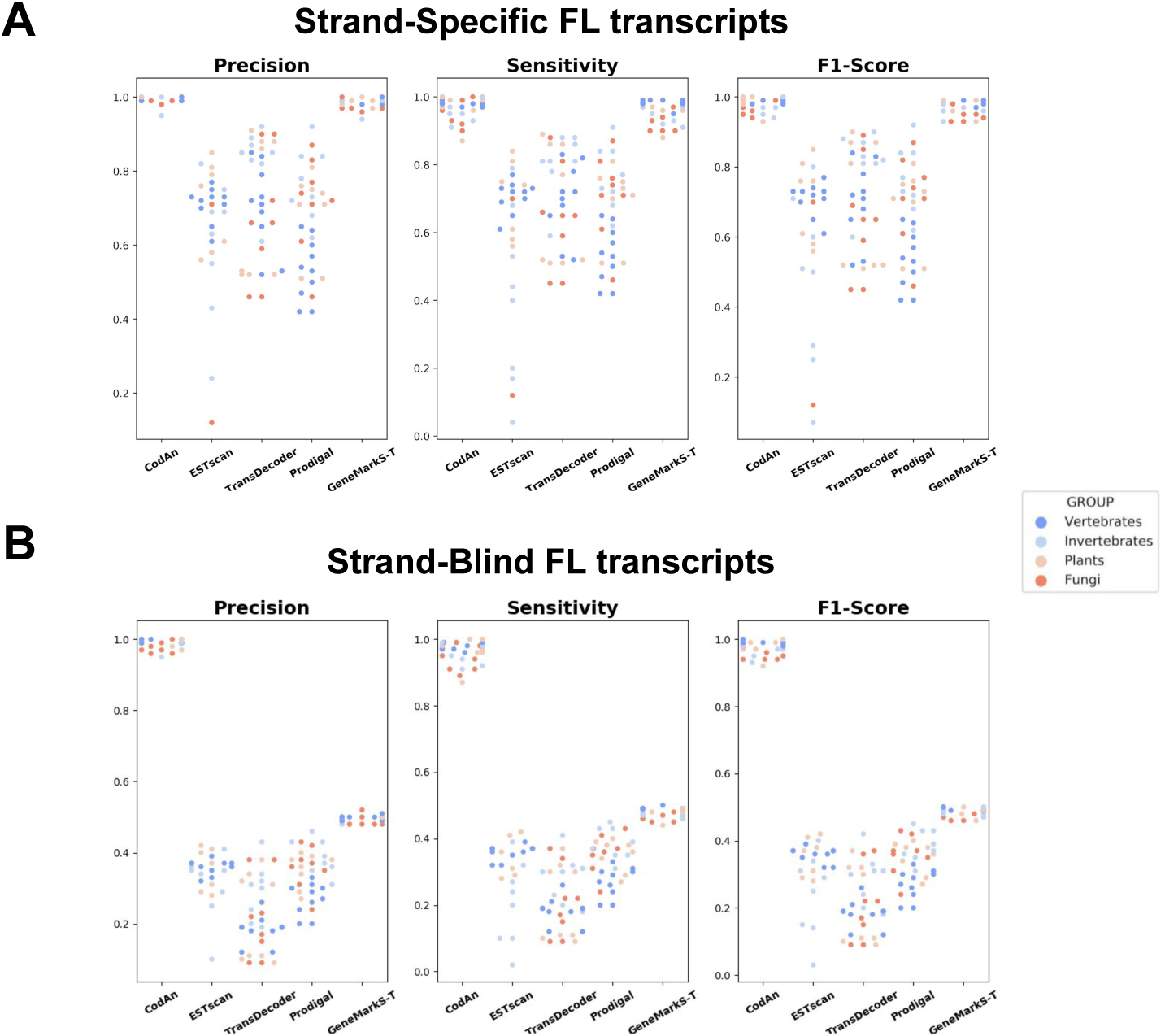
Scatter plot of the Full transcript test results. Each dot represents a different organism, coded by organism group. Results are grouped vertically by predictor. (A) Precision, Sensibility and F1-Score obtained by each tool in the strand-specific test; each dot represnts. (B) Precision, Sensibility and F1-Score obtained by each tool in the strand-blind test.

When considering strand-blind sets CodAn significantly outperforms all other applications in stop codon prediction (Table 3; Figure 1B), the F1-scores are at least 40% higher than other approaches, with consistently higher Precision and Recall values in all organisms (Table S2 C and S2 D at Supplemental Table S2). In fact, CodAn is the only software for which strand-specific and strand-blind results are almost the same, with F1-score values consistently over 95%. Considering that most RNA-seq projects perform sequencing with an unknown orientation of the transcript being sequenced, it is relevant to use predictors that present high precision independently the orientation of the CDS in the transcripts.

**Table 3:** Average and standard deviation of precision, Recall and F1-score obtained by each tool in the Strand-Blind full transcript sets analyzed. Bold font highlight the higher value for each group.

| Group | Predictor | Precision | Recall | F1-score |
| --- | --- | --- | --- | --- |
| Vertebrates | CodAn | **0.99 ± 0.00** | **0.98 ± 0.01** | **0.99 ± 0.01** |
| | ESTscan | 0.36 ± 0.02 | 0.36 ± 0.02 | 0.88 ± 0.13 |
| | TransDecoder | 0.18 ± 0.04 | 0.18 ± 0.04 | 0.93 ± 0.05 |
| | Prodigal | 0.27 ± 0.05 | 0.27 ± 0.05 | 0.93 ± 0.02 |
| | GeneMarkS-T | 0.50 ± 0.00 | 0.49 ± 0.00 | 0.49 ± 0.00 |
| Invertebrates | CodAn | **0.99 ± 0.02** | **0.95 ± 0.03** | **0.97 ± 0.02** |
| | ESTscan | 0.31 ± 0.10 | 0.21 ± 0.13 | 0.24 ± 0.12 |
| | TransDecoder | 0.29 ± 0.08 | 0.28 ± 0.07 | 0.29 ± 0.08 |
| | Prodigal | 0.39 ± 0.05 | 0.39 ± 0.05 | 0.39 ± 0.05 |
| | GeneMarkS-T | 0.49 ± 0.01 | 0.48 ± 0.01 | 0.48 ± 0.01 |
| Plants | CodAn | **0.99 ± 0.01** | **0.96 ± 0.04** | **0.98 ± 0.02** |
| | ESTscan | 0.35 ± 0.06 | 0.35 ± 0.06 | 0.35 ± 0.06 |
| | TransDecoder | 0.22 ± 0.13 | 0.21 ± 0.12 | 0.21 ± 0.12 |
| | Prodigal | 0.35 ± 0.05 | 0.35 ± 0.05 | 0.35 ± 0.05 |
| | GeneMarkS-T | 0.49 ± 0.01 | 0.48 ± 0.02 | 0.49 ± 0.01 |
| Fungi | CodAn | **0.98 ± 0.01** | **0.94 ± 0.04** | **0.96 ± 0.02** |
|  | ESTscan | NA | NA | NA |
| | TransDecoder | 0.21 ± 0.11 | 0.20 ± 0.10 | 0.21 ± 0.11 |
| | Prodigal | 0.36 ± 0.06 | 0.35 ± 0.06 | 0.36 ± 0.06 |
| | GeneMarkS-T | 0.49 ± 0.01 | 0.47 ± 0.02 | 0.48 ± 0.01 |

### Prediction accuracy assessment: experimentally validated strand-specific full transcripts

Next, we evaluated the accuracy for complete CDS prediction using a set of full transcripts of *H. sapiens, M. musculus, D. rerio, D. melanogaster* and *A. thaliana* with their respective start codons validated and annotated by Ribo-seq experiments [15, 17].

The tests revealed that GeneMarkS-T and CodAn presented higher performance than the other tools, but this time with a clear advantage for CodAn, with a higher percentage of correct predictions in 5 of the 7 benchmarks and small advantage in two (Table∼4, Figure 2; Table S2 E at Supplemental Table S2). CodAn presented higher rates of correct predictions and an almost perfect score for predicting the stop codon position (over 97% of the predictions in all datasets). These results confirm the consistent advantage of CodAn in full CDS prediction obtained in the first 34 benchmarks (Tables S2 A and S2 B in Supplemental Table S2).

**Table 4:** Precision, recall and F1-score for the prediction in datasets with start codons confirmed by riboseq experiments. True positives are sequences with the whole CDS predicted correctly (start and stop codon). Bold font highlight the higher value for each dataset.

| Species | Reference | Dataset size | Predictor | Precision | Recall | F1-score |
| --- | --- | --- | --- | --- | --- | --- |
| *H.sapiens* | Lim et al.. 2018 | 14193 | CodAn | **0.81** | **0.75** | **0.78** |
|  |  |  | ESTScan | 0.23 | 0.22 | 0.23 |
|  |  |  | TransDecoder | 0.37 | 0.35 | 0.36 |
|  |  |  | Prodigal | 0.27 | 0.27 | 0.27 |
|  |  |  | GeneMarkS-T | 0.67 | 0.63 | 0.65 |
| *H.sapiens* | Lee et al.. 2012 | 5727 | CodAn | **0.99** | **0.97** | **0.98** |
|  |  |  | ESTScan | 0.68 | 0.68 | 0.68 |
|  |  |  | TransDecoder | 0.60 | 0.59 | 0.60 |
|  |  |  | Prodigal | 0.49 | 0.49 | 0.49 |
|  |  |  | GeneMarkS-T | 0.98 | 0.96 | 0.97 |
| *M.musculus* | Lim et al.. 2018 | 20326 | CodAn | **0.85** | **0.83** | **0.84** |
|  |  |  | ESTScan | 0.30 | 0.30 | 0.30 |
|  |  |  | TransDecoder | 0.39 | 0.38 | 0.39 |
|  |  |  | Prodigal | 0.29 | 0.29 | 0.29 |
|  |  |  | GeneMarkS-T | 0.70 | 0.68 | 0.69 |
| *M.musculus* | Lee et al.. 2012 | 2701 | CodAn | **1.00** | **0.98** | **0.99** |
|  |  |  | ESTScan | 0.74 | 0.74 | 0.74 |
|  |  |  | TransDecoder | 0.67 | 0.65 | 0.66 |
|  |  |  | Prodigal | 0.52 | 0.52 | 0.52 |
|  |  |  | GeneMarkS-T | 0.97 | 0.95 | 0.97 |
| *D.rerio* | Lim et al.. 2018 | 13954 | CodAn | **0.86** | **0.85** | **0.86** |
|  |  |  | ESTScan | 0.44 | 0.44 | 0.44 |
|  |  |  | TransDecoder | 0.67 | 0.65 | 0.66 |
|  |  |  | Prodigal | 0.47 | 0.47 | 0.47 |
|  |  |  | GeneMarkS-T | 0.80 | 0.78 | 0.79 |
| *D.melanogaster* | Lim et al.. 2018 | 13653 | CodAn | **0.90** | **0.90** | **0.90** |
|  |  |  | ESTScan | 0.68 | 0.66 | 0.67 |
|  |  |  | TransDecoder | 0.57 | 0.56 | 0.56 |
|  |  |  | Prodigal | 0.56 | 0.56 | 0.56 |
|  |  |  | GeneMarkS-T | 0.86 | 0.83 | 0.85 |
| *A.thaliana* | Lim et al.. 2018 | 6947 | CodAn | **0.87** | **0.85** | **0.86** |
|  |  |  | ESTScan | 0.59 | 0.58 | 0.58 |
|  |  |  | TransDecoder | 0.67 | 0.65 | 0.66 |
|  |  |  | Prodigal | 0.56 | 0.56 | 0.56 |
|  |  |  | GeneMarkS-T | 0.82 | 0.79 | 0.80 |

**Figure 2:**
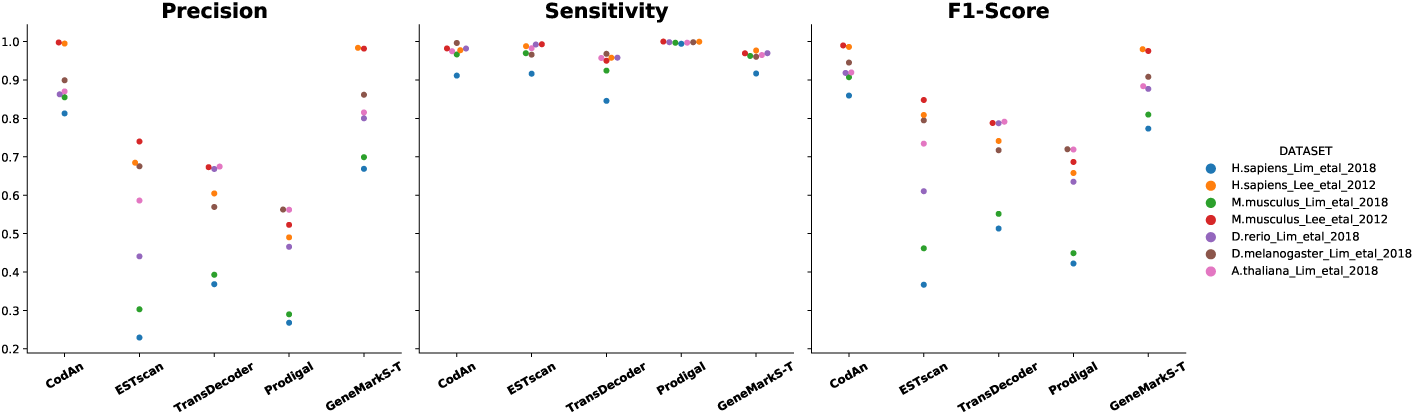
Scatter plot of the precision, sensitivity and F1-score obtained by each tool in the Ribo-seq experimentally validated datasets considering the full CDS region.

In summary, for full transcripts CodAn significantly outperforms other available software in predicting CDS for full transcript of unknown orientation, and increases precision in full CDS prediction, while still matching the best stop codon precision measurements. This indicates that CodAn it the best choice when the annotation of the whole coding sequence is necessary for the subsequent analysis.

### Prediction accuracy assessment: partial transcripts

In most real-life situations, transcript sequencing programs produce initially a high rate of partial transcripts. It is therefore relevant to measure the accuracy of predictions also for these sequences. To form a more precise picture we separately measure the accuracy for prediction in transcripts consisting of: (i) only CDS nucleotides; (ii) 5’UTR and CDS nucleotides, (iii) CDS and 3’UTR nucleotides. These data sets presented a much harder challenge for all of the applications used in the comparison Figure 3; Table S2 F in Supplemental Table S2). For the NoStart dataset, only CodAn was able to correctly identify a significant number of CDSs, in this case with very good results, averaging over 97% in F1 score values. The situation changed for the NoStop datasets, but still with clear advantage for CodAn: average F1 score for CodAn was above 58%, in comparison to a maximum of 27% for the other applications. Finally, for the NoStartNoStop (CDS only) sequences, CodAn F1 scores were, again, averaged more than 96%, while F1 scores for the other applications was always below 13%.

**Figure 3:**
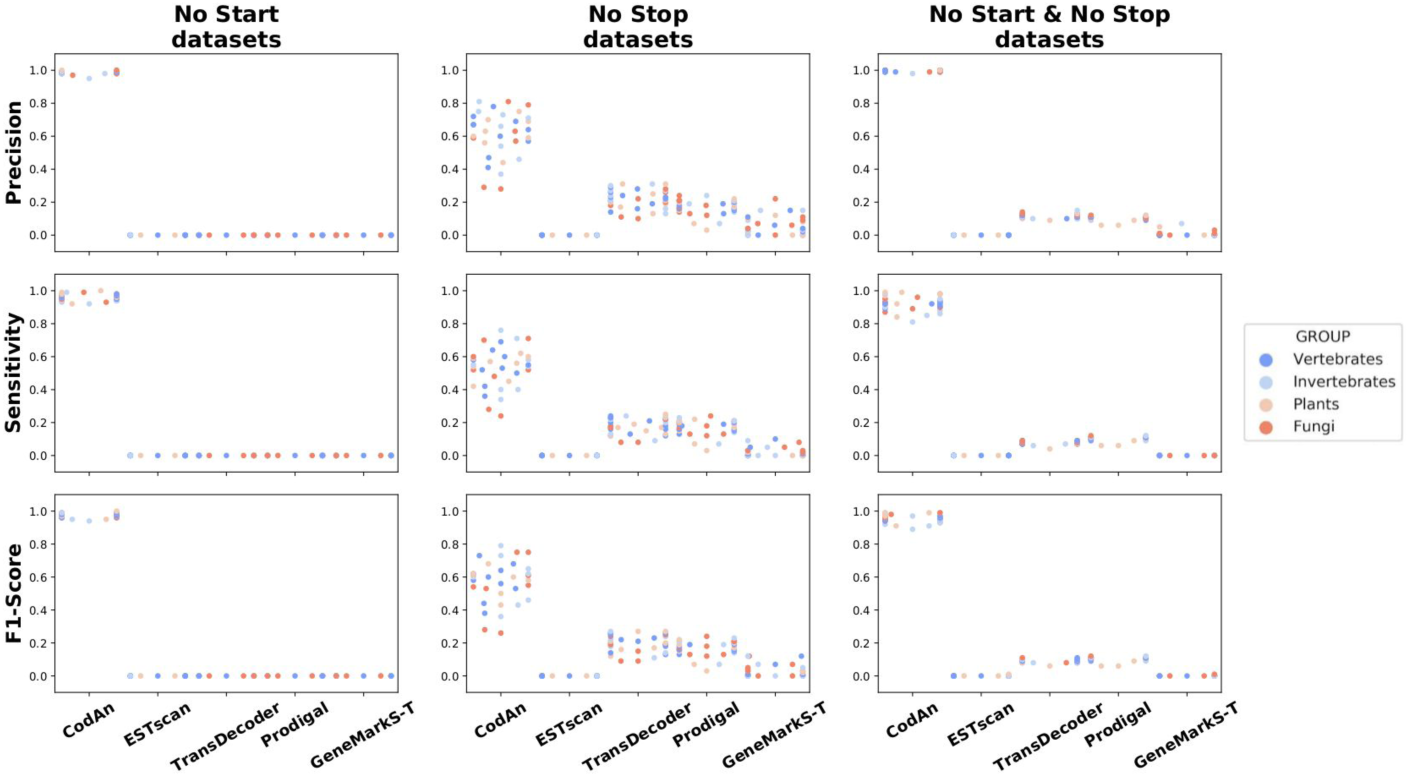
Scatter plot of the partial transcript test results. Each dot represents a different organism, coded by organism group. Results are grouped vertically by predictor. From right to left the plots represent the Spider plots showing the Percentage of correct predictions on “No Start”, “No Stop” and “No Start & No Stop” tests performed on all species analyzed in the present study.

These results showed that other approaches fail to obtain even modest precision or recall rates for the prediction of CDSs in partial transcripts, with CodAn achiving consistently higher rates. This clearly indicte CodAn as the best approach to handle cases where the partial transcripts are highly abundant in the transcriptome assembly. In fact, it is a common feature in *de novo* assemblies, which partial transcripts can represent up to 50% of the sequences assembled [8].

### False-positives assessment using partial 3’UTR and ncRNA transcripts

Different sequencing protocols can produce two types of negative sequences when considering CDS prediction: UTR-only sequences or ncRNA sequences.

To estimate the rate in which such transcripts have false-positive predictions in the first case we ran all applications in the 3’UTR sequences of the previous data sets. The results show that CodAn and TransDecoder, as a rule, presented the lowest number of false-positives, whereas Prodigal presented the highest number of false-positives (Figure 4A; Table S2 G at Supplemental Table S2). TransDecoder presented the best overall performance with better average specificity values for Invertebrates (95% vs 90%), Plants (95% vs 90%) and Fungi(95% vs 90%). The only exception was for Vertebrates with GeneMarkS-T presenting a Specificity of 97%, against 95% of CodAn and 90% of TransDecoder.

**Figure 4:**
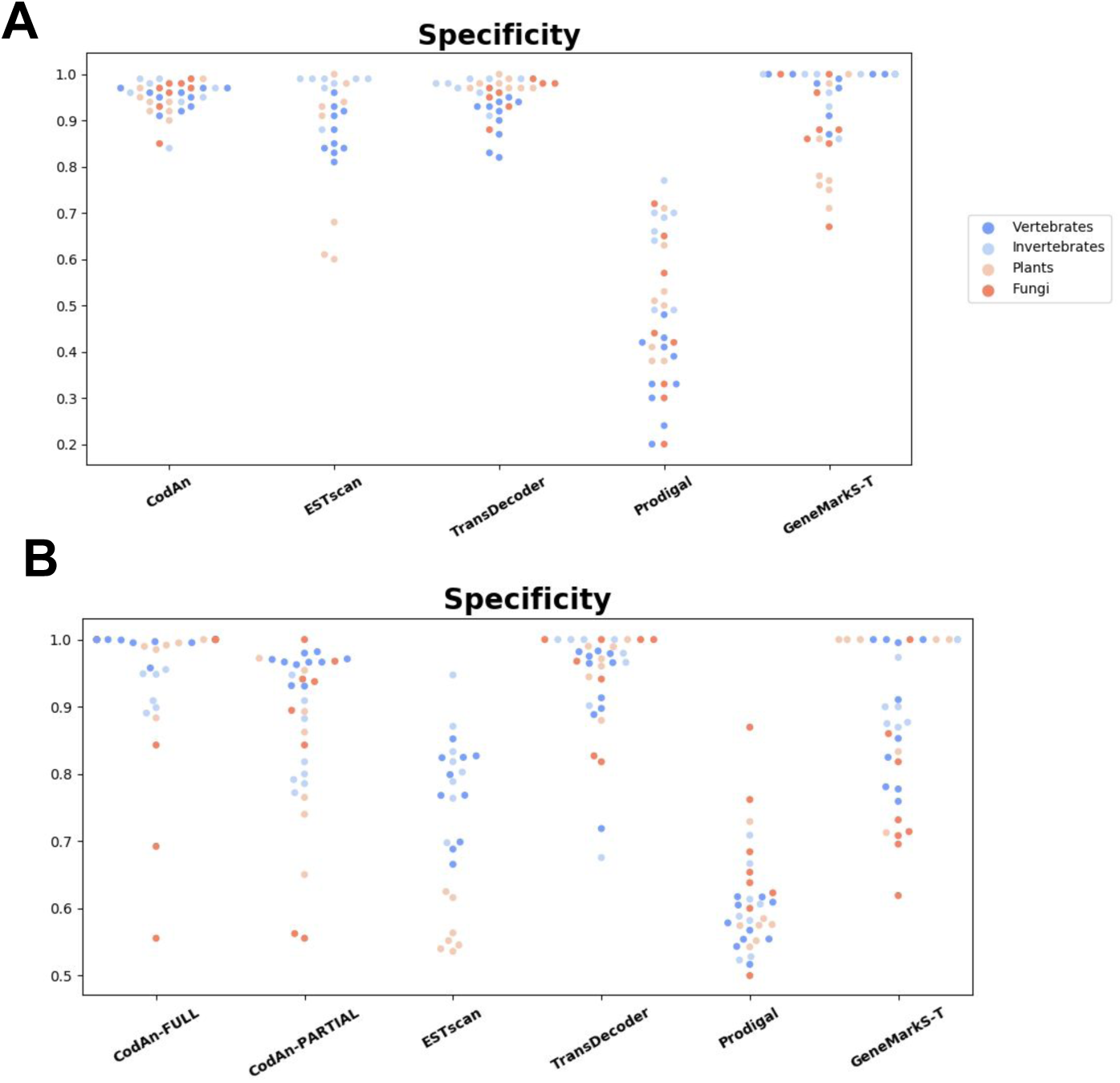
Scatter plot of Specificity obtained by each tool in the prediction. Each dot represents a differnt organism, color-coded by organism group. Results are grouped vertically by predictor. We used two negative datasets (A) 3’UTR region datasets and (B) Full ncRNA datasets. Each dot represents a differnt organism, color-coded by organism group.Results are grouped vertically by predictor.

Specificity assessment for ncRNA sequences showed that the full transcript model of CodAn presents the best performance of all predictors. If, instead, we use the partial model of CodAn, its performance is higher for Vertebrates, whereas Transdecoder presented a slightly better performance in the other groups (Figure 4B; Supplementary Table S2 H at Supplemental Table S2). Overall, both models of CodAn presented satisfactory results on specificity tests.

## Conclusions

We presented CodAn, a software that generates highly confident transcript characterization in Eukaryote species in all common sequencing project situations. This high confidence is achieved by the use of multiple probabilistic models integrated using a Generalized Hidden Markov Model (GHMM). The design of CodAn was based on the development of model parameters for four groups of Eukaryotes: Vertebrates, Invertebrates, Plands and Fungui. Each parameter set was estimated based on a mix of reference transcripts from several species of one of the organism groups. CodAn can run in any Desktops/Laptops or take advantage of large multi-processor servers based on UNIX OS.

We showed that these generic models work well and result in a reliable characterization of transcripts on a wide range of Eukaryote species. Considering the benchmarks used in the present analysis, CodAn had a clear performance advantage when considering all common RNA sequecing projects situations, in particular with strand-blind full sequences and partial sequences.

In summary, our data indicates that CodAn is the best approach to be applied on studies focusing to characterize the CDS regions and the UTR landscape of partial and/or full transcripts, an can help the improvement of current and future gene annotation for transcriptomes of Eukaryote species, whic is a field under constant expansion [24].

## Methods

### Algorithm implementation

CodAn uses two different architectures for analysing transcripts, one for full and one for partial transcripts (Figure 5. Both architectures are described using Generalized Hidden Markov Models implemented using the ToPS probabilistic framewok [12]. Of note, we partition our probabilistic model in GG content specific sub-models [28]. More details on the probabilistic model are described in the supplementary file Supplemental Methods.

**Figure 5:**
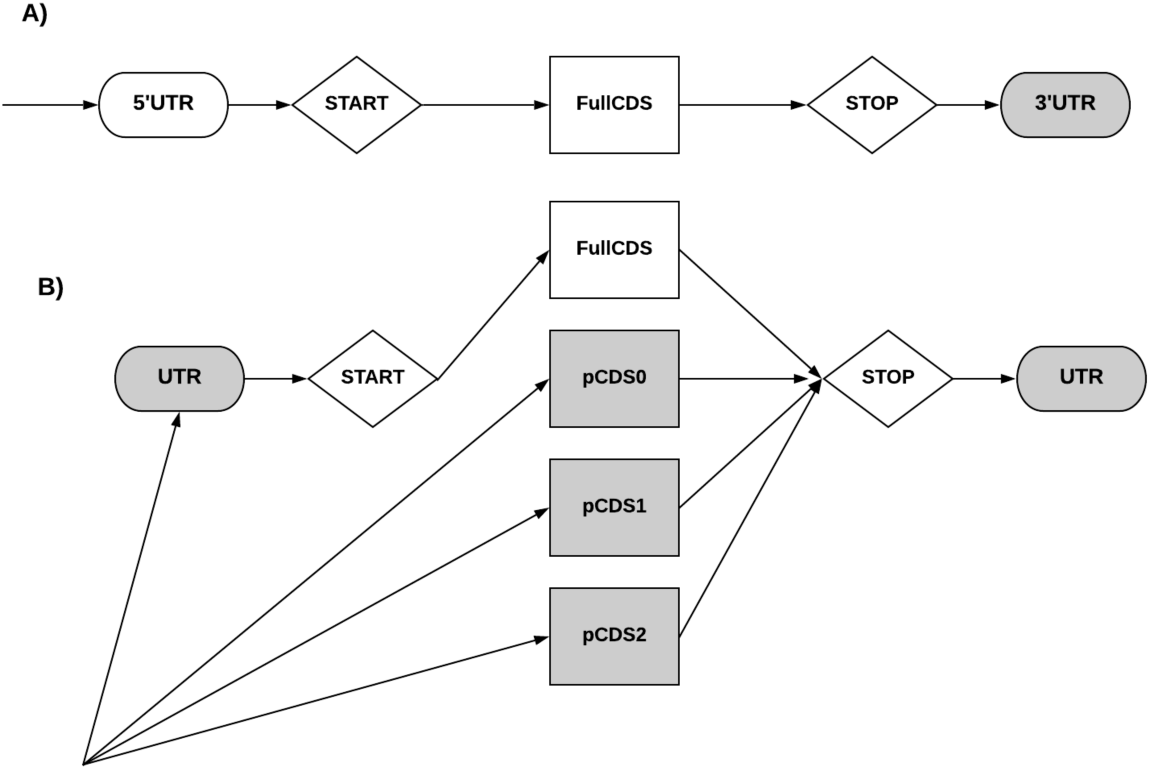
The two GHMMs representing transcripts. A) Full Transcript model, grey figures represent final states, the arrows represent the flow of the architecture, indicating only one initial state, 5’UTR. B) Parcial transcript model, grey figures represent final states, the arrows represent the flow of the architecture. The four states represented by circles are states with explicit duration distribution that emit the protein-coding region: fullCDS, pCDS0, pCDS1, and pCDS2. The state fullCDS models a complete coding region. The states pCDS0, pCDS1, pCDS2 represent partial coding regions that start, respectively, at frame 0, 1, and 2. The state labeled UTR can be used to represent either the 5’UTR or the 3’UTR. The 3’UTR state represents 3’UTRs. The states *Start* and *Stop* in diamonds have a fixed length duration, and they represent the start codon and stop codon, respectively.

CodAn uses ToPS [12] to implement the Generalized Hidden Markov Model architectures, Python (v.3.6.8) and Perl (v5.26.1) scripts to prepare and process data for the ToPS probabilistic framework.

For each architecture four different sets of parameters were estimated, corresponding to four organism groups: Vertebrates, Invertebrates, Plants and Fungi. By default, CodAn takes as input transcripts in FASTA format, performs the prediction and returns three FASTA files, containing the CDS, 3’UTR and 5’UTR sequences predicted for each transcript, and a GTF file, containing the annotations of the predictions for each transcript.

### Training Sets

CodAn uses probabilistic models for which we need to estimate the parameters. For this, we used training sets with reference sequences from different species downloaded from the RefSeq Database at NCBI (release number 94; ftp://ftp.ncbi.nlm.nih.gov/refseq/). Due to lack of complete annotations of transcripts for *C. elegans* at RefSeq, we used the sequences deposited at the WormBase (release WS270; ftp://ftp.wormbase.org/pub/wormbase/). We retrieved sequences following three criteria: (1) presence of a reviewed and/or curated status; (2) validated expression status; and (3) full-length transcripts. We estimated 4 different parameter sets, each one targeted to a different group of Eukaryotic organisms: vertebrates, invertebrates, plants, and fungi. The training sets for each parameter set contained reference transcript sequences of a mix of species from each group (detailed in Supplementary Table S1 A at Supplemental Table§1).

### Comparison protocol

We compared the prediction performance of CodAn against that of ESTscan (v3.0.3; [19]), TransDecoder (v5.5.0; [6]), Prodigal (v2.6.3; [10]) and GeneMarkS-T (v5.1; [29]). We used all tools with default parameters following their usage guidelines, as the fine-tuning of parameters of each tool are beyond the scope of this analysis. For ESTscan we used the pre-trained models either of the species being tested or the closest related species when the species-specific model was not available (the pre-trained models used for each species are specified in Table S1 B at Supplemental Table S1). Since there was no Fungi model for ESTScan, we did not perform comparison tests for this tool in the Fungi group. For Prodigal, we used the mode directed to predict intron-less genes (“switched-off RBS model”), which can be applied to predict coding regions in transcripts of Eukaryotes.

We performed a comparison in both full transcript and partial transcript sets. Following Tang and collaborators [29], we used both annotated and Ribo-seq validated full transcripts. The first for evaluating accuracy of stop codon prediction, the second for evaluating full CDS prediction accuracy. In all tests we measured the Precision (computed as TruePositives / (TruePositives + FalsePositives), Recall (computed as TruePositives / (TruePositives + FalseNegatives)) and F1-score (computed as 2 * (Precision * Recall) / (Precision + Recall)). Following [29], we considered True Positives as the predictions that exactly matched the reference annotation, False Positives as the predictions that presented any difference from the reference annotation, and False Negatives as the sequences with no predictions. In this sense, for the annotated full transcripts we considered as True Positives the predictions that correctly matched the annotated stop codon, and false positives all other predictions. For both the Ribo-seq validated sequences and for the partial sequences we considered True Positives all predictions that correctly matched the whole CDS of the transcripts.

We adopted the most common interpretation of the concepts of True Positive, False Positive and False Negatives used in gene prediction. These measures would be sufficient in the ideal situation where all sequences are mRNA transcripts with a CDS region. However, in transcriptomic projects sequences with no CDS region can be present, either being just UTRs or, depending on sequencing protocol, ncRNAs. To evaluate the rate of false discoveries, we also compared Specificity (computed as FalsePositives / (FalsePositives + TrueNegatives)) of the various approaches. For this, we used two different negative datasets: 3’UTR regions and ncRNAs. It is important to note that here the definition of False Positives is somewhat different from that used in computing Precision, Recall, and F1-score.

### Testing sets

The test sets for comparison against other approaches consisted of transcript data from 34 Eukaryote species that are of interest in the fields of evolutionary and biomedical studies and/or highly used in food production (Table 1). For each of the 34 organisms, we retrieved 2000 randomly selected full transcripts presenting the following three criteria: (1) validated expression status; (2) full length; and (3) full CDS annotation. None of these sequences included any of the transcripts used for training the probabilistic model. For each transcript set, we generated 7 distinct validation sets: two sets with full transcripts, four sets with partial transcripts and one set with ncRNA sequences.

The first full transcript set (Full Strand-Specific) included all 2000 transcripts as downloaded from the database. For the second, full transcript set (Full Strand-Blind), intended to measure the performance of the predictors in sequences with unknown translation direction, we randomly selected half of the sequences in the previous datasets (1000 transcripts) and replaced it for its reverse complement.

To compare the prediction of complete CDSs, we followed the approach of Tang and collaborators [29], using a set of full transcripts with their respective start codons validated and annotated by Ribo-seq experiments [15]. We, however, extended the number of species in the validation including *H. sapiens, M. musculus, D. rerio, D. melanogaster* and *A. thaliana* [17]. For the data previously analyzed by [15], we selected transcripts where the annotated start codon at RefSeq matched to the start codon confirmed by the Ribo-seq data resulting in 5727 and 2701 sequences for *H. sapiens* and *M. musculus*, respectively. On the data analyzed by [17], we considered only the full transcripts with the curated annotation by the Ribo-seq data, which led to 14193, 20326, 13954, 13653 and 6947 sequences for *H. sapiens, M. musculus, D. rerio, D. melanogaster* and *A. thaliana*, respectively.

For the partial transcripts datasets, we considered that the *de novo* assemblies can result in partial transcripts with no start codon and/or no stop codons. For the first partial transcript set (No Start), we randomly selected, for each transcript of the full transcript data set, a cutting point in the CDS region and pruned the 5’ part, eliminating the start codon. In the second partial transcript set (No Stop), we randomly selected, for each transcript a new cutting point in the CDS region and pruned the 3’ part, eliminating the stop codon. In the third partial transcript set (No Start & No Stop), we randomly selected, for each transcript two cutting points and eliminated the 5’ and the 3’ ends of the transcript, retaining only part of the CDS region. Cutting points were selected to guarantee a minimum size of 150nt for the resulting sequences. In cases where the whole transcript was smaller than 150nt we only pruned the sequence at the start and/or stop codon, depending on the dataset.

We used two different data sets to evaluate Specificity: 3’UTR sequences and ncRNA sequences. The 3’UTR data set consisted of the complete 3’UTR regions of each transcript in the original full transcript dataset. The 3’UTR set was designed to be a realistic negative set for protein-coding transcriptome projects, when the experimental design leads to a selection of mRNA transcripts based on poly-A selection. Additionally, to test the specificity in RNASeq projects with no poly-a specificity we used ncRNA sequences. For this we created a data set containing all ncRNA sequences longer than 200nt length available for each species in the RFAM database (release 14.1;https://rfam.xfam.org/). To make the specificity test more close to a real transcriptome assembly and fair for the self-training algorithms, we used a mix of sequences containing a proportion of 500 sequences of full-length transcripts and 500 sequences of partial transcripts within the ncRNA sequences.

## Declarations

### Author’s contributions

PGN designed the initial application, implemented the python scripts, designed the train and test datasets, designed and ran the tests, designed automated scripts to analyze the data and plot the charts and wrote the article. AYK designed and implemented the ToPS C++ probabilistic models, designed the probabilisic model architecture and wrote the article. AMD coordinated the implementation of all parts of the project, supervised the design of the test in procedures and wrote the article.

### Availability of Data and Materials

The datasets used in the present study, along with the code for the CodAn software, are available in the github repository: https://github.com/pedronachtigall/CodAn.

## Supplementary Files

**Supplemental_File_1**

Supplementary data in pdf format.

**Supplemental_Table_S1**

Supplementary data in xls format.

**Supplementary file 3**

Supplementary data in xls format.

