## Supplemental Methods for "CodAn: predictive models for the characterization of mRNA transcripts in Eukaryotes"

### PROBABILISTIC MODEL

The 8 states of the architecture are implemented in ToPS (Kashiwabara et al., 2013). The global architecture is implemented using a Generalized Hidden Markov Model. The emission and the duration models used for each state varies depending on the region being represented, as detailed below.

The states CDS0, pCDS1, pCDS2 represent regions coding partial proteins that start at frame 0, 1, and 2, respectively. The state full CDS models a coding region of a complete protein. Each one of these protein-coding regions is represented using a three-periodic Interpolated Markov Model (Salzberg et al., 1998) of order 4. All these states have an explicit duration model with distribution based on the training set.

The Start Stop states represent the start codon region and the stop codon region, respectively. The Start state represents a sequence of length 25nt that we can break it in three segments: (i) a sequence segment with 20 positions representing the region before the ATG; (ii) a sequence with three positions representing the ATG sequence itself; (iii) and a sequence with four positions represent the region after the ATG. We used a windowed weight array model (Burge and Karlin, 1997) of order 1 and vicinity length 4 for the first segment and a Weight Array Model (Zhang and Marr, 1993) of order 4 for the third segment. The Stop state represents sequences of three nucleotides with the pattenns TAA, TAG, or TGA. Probabilities of each of the patterns correspond to their frequency in the training set.

The 5'UTR and 3'UTR states represent the 5' and 3' untranslated regions, respectively. They are represented using an Interpolated Markov Model of order 4 (Salzberg et al., 1998). The state duration is represented with a geometric length distribution with parameter estimated on the duration of the 5'UTR and 3'UTR sequences of the training set.
